# Photosystem genes in chloroplast and their interacting partners: A case for molecular adaptation to carnivory in *Nepenthaceae*

**DOI:** 10.1101/2022.05.24.493228

**Authors:** Neeraja M Krishnan, Binay Panda

## Abstract

Molecular adaptations are prevalent in carnivorous plants in response to habitat and environmental stress. We used the chloroplast genome and characterized the specific adaptations in the photosystem genes and their interacting partners in *Nepenthes khasiana*, a carnivorous pitcher plant. When compared with the carnivorous and non-carnivorous groups across Caryophyllales, Lamiales, Poales, Ericales, and Oxalidales, we found *Nepenthes*-specific changes in psaA, psaB, psaC and psaH. Of these, only a single amino acid change each, G147 in the protein psaA and R40 in the protein ndhD, impacted the three-dimensional structural conformation of the corresponding proteins. Modeling the interaction between the psaA and the ndhD proteins identified group-specific changes between the models between *Nepentheceae* versus others. The least distance between the structure-impacting residues of psaA and ndhD was 25.9 Å for *Nepenthes* and 19.4 Å for non-*Nepenthes* models. Given that the chloroplast ndh and photosystem I subunits form a large super-complex with the light-harvesting carrier proteins from the nucleus to mediate cyclic electron transport, our observations may indicate specific adaptations in the cyclic electron transport arm of the photosynthetic machinery in the *Nepenthes* species.

## INTRODUCTION

Carnivorous plants bear both plant and animal traits and typically grow on nutrient-poor soil or in dense habitats impermeable to sunlight. They passively prey on unsuspecting insects and small animals and digest nutrients from them with the help of modified tactile plant parts that function as traps. In carnivorous plants, the complex modifications represent sophisticated adaptations at structural and functional levels where the benefits surpass the metabolic costs (Adamec, 2002, Plachno et al., 2020, Thorogood and Bauer, 2020). Members of the carnivorous plant group *Nepenthaceae*, a monotypic family in the order Caryophyllales, are spread throughout the old-world tropics except for *Nepenthes khasiana*. The group has 120 species of pitcher plants that grow in hot and humid areas with large modified leaves turned pitchers in nutrient-poor soil. Previous works suggested that *Nepenthaceae* group members originated from an ancient stock before moving to Indo-China and the Malay Archipelago (Biswal et al., 2018, Meimberg et al., 2006, Meimberg et al., 2001). Members of the group *Nepenthaceae* are a perfect example where heterotrophic adaptations are prevalent. In these plants, the animal dependence is only to supplement the nutrient requirements of the plant and not to do away with the photosynthetic machinery all together (Behie and Bidochka, 2013, Chou et al., 2014, Gorb and Gorb, 2006, Gray et al., 2017, Heubl et al., 2006, Krause, 2008, Meimberg and Heubl, 2006, Moran et al., 2013, Murphy et al., 2020, Owen and Lennon, 1999, Shen et al., 2021).

The photosynthetic genes encoded by chloroplasts in some carnivorous plants undergo a selection process that is comparatively relaxed with a skew in favor of non-synonymous to synonymous substitution rate ratios (Krause, 2008). This is relatively understudied in the group *Nepentheceae*. We obtained the sequence of the chloroplast genome of *Nepenthes khasiana* from GenBank (Accession NC_051455.1) in order to address specific changes in the photosystem genes coded by the chloroplast and their interacting partners. By comparing the photosystem gene subunits across species from five carnivorous plant orders, Caryophyllales, Lamiales, Ericales, Poales and Oxalidales, and eleven non-carnivorous control species in Caryophyllales and non-Caryophyllales, we analyzed the structural and functional impact of the changes unique to the *Nepenthaceae* group. We performed similar analyses in ndh genes, which interact with the photosystem genes during cyclic electron transport process (Endo et al., 2008, Florez-Sarasa et al., 2016, Ishikawa et al., 2016, Joet et al., 2002, Kato et al., 2021, Kouril et al., 2014, Laughlin et al., 2019, Pan et al., 2020, Peng et al., 2011, Shikanai, 2014, Shikanai, 2016, Storti et al., 2020a, Storti et al., 2020b, Strand et al., 2017, Yin and Struik, 2018, Zhao et al., 2020).

We then studied the impact of *Nepenthaceae*-specific structural changes in *psaA* and *ndhD* on the interaction between these genes and observed *Nepenthaceae*-specific changes in *psaA* and *ndhD* to also impact interactions between them. The photosystem I (PSI) sub complex is known to form a super-complex with the NDH sub-complex along with other helper genes during the cyclic electron transport process (Shen et al., 2021, Yamori et al., 2016). This super complex stabilizes ndh and facilitates in achieving the necessary amount of ATP homeostasis required during the carnivorous aspects of prey feeding and retention (Pavlovic et al., 2010, Maurer et al., 2020, Shikania 2016). Piecing together these evidences from various carnivorous plants and study of chloroplast ndh in general (Shikania 2016, Peng et al., 2011), we postulate in this paper that the *Nepenthaceae*-specific structure impacting changes in *psaA, ndhD* and *psaA*—*ndhD* interactions most likely point to adaptations facilitating the cyclic electron transport portion of the photosynthetic machinery, specific to this group.

## MATERIALS AND METHODS

### Chloroplast genomes used for comparison

The sequence and the assembly information on the chloroplast genome of *Nepenthes khasiana* was obtained from NCBI (NC_051455.1). Both the carnivorous and non-carnivorous species, where complete chloroplast genomes were available, were considered for analyses. **Table 1** lists all the species used for the analysis. The orders where carnivory evolved independently (Ellison and Gotelli, 2009), namely, Caryophyllales (*Nepenthaceae* (including *Nepenthes khasiana -* NC_051455.1), *Drosophyllaceae, Dioncophyllaceae* and *Droseraceae*), Lamiales (Lentibulariaceae, Ericales, Poales) and Oxalidales (*Cephalotus*) were considered for the comparative analyses. Non-carnivorous plants from these orders and various other non-carnivorous species including *Arabidopsis thaliana, Ginkgo biloba, Epipactis helleborine, Dendrobium nobile, Neottia nidus-avis, Aneura mirabilis, Pogonia japonica, Monotropa uniflora, Cuscuta reflexa, Physcomitrium patens* and *Alsophila gigantea* were included as controls.

**Table 1.** List of species with available chloroplast genomes used for the study.

| Carnivorous plant orders |  |  |  |  |  |  |  |  |  |  |
| --- | --- | --- | --- | --- | --- | --- | --- | --- | --- | --- |
| Caryophyllales |  |  | Lamiales |  | Oxalidales |  | Poales |  | Ericales |  |
| Non-carnivorous plant controls |  |  |  |  |  |  |  |  |  |  |
|  |  | Mycoheterotrophic plants |  |  |  |  | Parasitic plants |  |  |  |
| Angiosperm | Gymnosperm | Non photosynthetic orchids | Photosynthetic orchids | Parasitic orchids | Non photosynthetic liverworts | Bog orchids | Holo-parasitic | Hemi-parasitic | Mosses | Ferns |
| *Arabidopsis thaliana* | *Ginkgo biloba* | *Epipactis helleborine* | *Dendrobium nobile* | *Neottia nidus-avis* | *Aneura mirabilis* | *Pogonia japonica* | *Monotropa uniflora* | *Cuscuta reflexa* | *Physcomitrium patens* | *Alsophila gigantea* |

### Comparative analyses of Nepenthaceae-specific amino acid changes in photosystem and ndh genes

Amino acid sequences for chloroplast-encoded genes were downloaded in the FASTA format for all Caryophyllales as of November 2021 from the NCBI protein database (https://www.ncbi.nlm.nih.gov/protein). The sequences were further converted to a 2 line FASTA format using the online DNA/RNA sequence conversion tool by bugaco.com (http://sequenceconversion.bugaco.com/converter/biology/sequences/). They were then parsed through, using the gene name bearing fasta headers using LibreOffice Calc’s Text-to-Columns on ‘.’, ‘[‘ and ‘]’ delimiters to extract sequences pertaining to photosystem I (psa), photosystem II (psb) and NADH dehydrogenase (ndh) subunits.

The amino acid sequences for photosystem and ndh subunits from *Nepenthes khasiana* were matched against the corresponding subunit sequences from other species using blastp (Altschul et al., 1990); https://blast.ncbi.nlm.nih.gov/Blast.cgi?PAGE=Proteins&PROGRAM=blastp&BLAST_PROGRAMS=blastp&PAGE_TYPE=BlastSearch&BLAST_SPEC=blast2seq&DATABASE=n/a&QUERY=&SUBJECTS=) with Align 2 or more sequences option, Max target sequences of 5000 and a word size of 2. From the BLAST hits, *Nepenthaceae*-specific changes were recorded.

The structural impact of mutations was assessed using the Missense3D server (http://missense3d.bc.ic.ac.uk/missense3d/;40) for all the *Nepenthaceae*-specific changes in photosystem and ndh proteins.

### Extending comparison of Nepenthaceae-specific structure-impacting changes across other carnivorous and non-carnivorous plants

We extended the comparison of *Nepenthaceae*-specific changes in psaA and ndhD that were structure-impacting to other carnivorous and non-carnivorous plants listed in **Table 1**.

### Modelling protein-protein interaction between psaA and ndhD

The GRAMM-X Protein-Protein Docking Web Server v.1.2.0 (http://vakser.compbio.ku.edu/resources/gramm/grammx; (Tovchigrechko and Vakser, 2006) was used to compute 300 models each of interactions between *psaA* and *ndhD* in *Nepenthes* and other species used for comparison. The models were individually visualized using iCn3D, a web-based structure viewer (https://www.ncbi.nlm.nih.gov/Structure/icn3d/full.html; (Wang et al., 2020) and the distance between 147 residue in *psaA* and 40th residue in *ndhD* were computed for these models. The distributions of these distances were then plotted as a frequency polygon using the Easy Frequency Polygon Maker tool provided by socscistatistics.com (https://www.socscistatistics.com/descriptive/polygon/default.aspx).

## RESULTS

### Nepenthaceae-specific changes across Caryophyllales in photosystem genes

We used the RefSeq curated sequences for Caryophyllales in the NCBI protein database. We used, out of a total of 84,732 chloroplast protein sequences, 3,392 sequences pertaining to psa subunits (698 *psaA*, 654 *psaB*, 651 *psaC*, 671 *psaI* and 718 *psaJ*) and 14,287 pertained to psb subunits (3,058 *psbA*, 977 *psbB*, 662 *psbC*, 876 *psbD*, 698 *psbE*, 653 *psbF*, 732 *psbH*, 664 *psbI*, 1,228 *psbJ*, 710 *psbK*, 637 *psbL*, 880 *psbM*, 754 *psbN*, 965 *psbT* and 793 *psbZ*) that belonged to the Caryophyllales order (**Appendix S1**).

Only *psaA, psbB, psbC* and *psbH* gene subunits were found to harbor *Nepenthaceae*-specific amino acid changes. These changes were G147A in *psaA*, N72T and V199A in *psbB*, Q39E in *psbC* and R30K in *psbH*. Of these, only G147A in *psaA* was found to have an impact on the structure (**Figure 1**). The GLY to ALA difference represents a replacement of buried residue with a Relative Solvent Accessibility (RSA) of 3.5% with a buried residue with a higher RSA of 4.7%.

**Figure 1.**
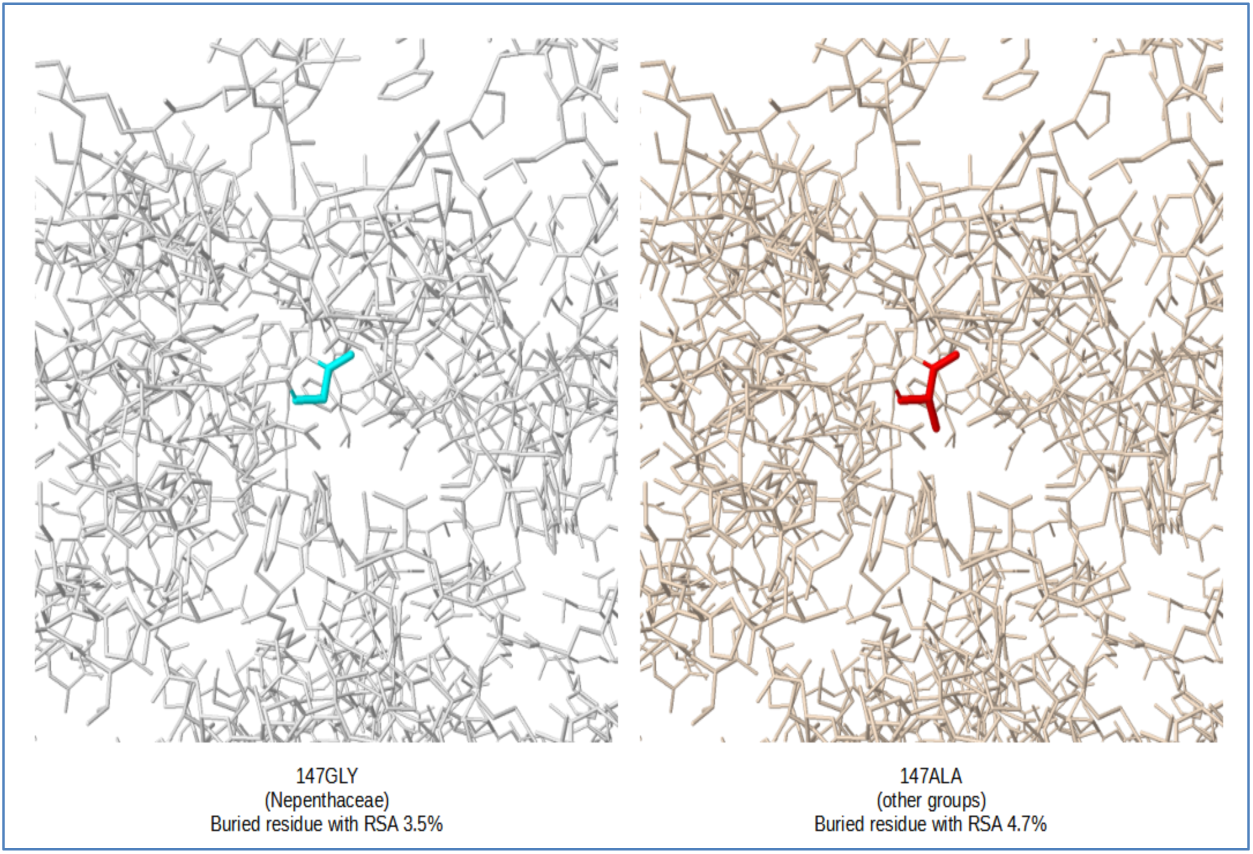
Characterization of structural impact of *Nepenthaceae*-specific amino acid changes in psaA.

### Nepenthaceae-specific changes across Caryophyllales in ndh genes

Out of the total of 84,732 chloroplast protein sequences for Caryophyllales in the NCBI protein database, 16,838 pertained to ndh subunits (606 *ndhA*, 1,232 *ndhB*, 633 *ndhC*, 587 *ndhD*, 608 *ndhE*, 1,624 *ndhF*, 606 *ndhG*, 603 *ndhH*, 606 *ndhI*, 662 *ndhJ* and 600 *ndhK*) that truly belonged to the Caryophyllales order (**Appendix S2**). Only *ndhD* gene subunit was found to harbor a structure impacting *Nepenthaceae*-specific amino acid change (**Figure 2**). This change was at residue 40 where the *Nepenthes* group alone has ARG as a buried charged residue, and is replaced by an uncharged residue in all other groups. The uncharged residue at this position varies between TRP for most Caryophyllales and non-carnivorous controls, LEU for some Basellaceae, some Droseraceae, and all Ancistrocladaceae among Caryophyllales and Lamiales, and SER for some Molluginaceae.

**Figure 2.**
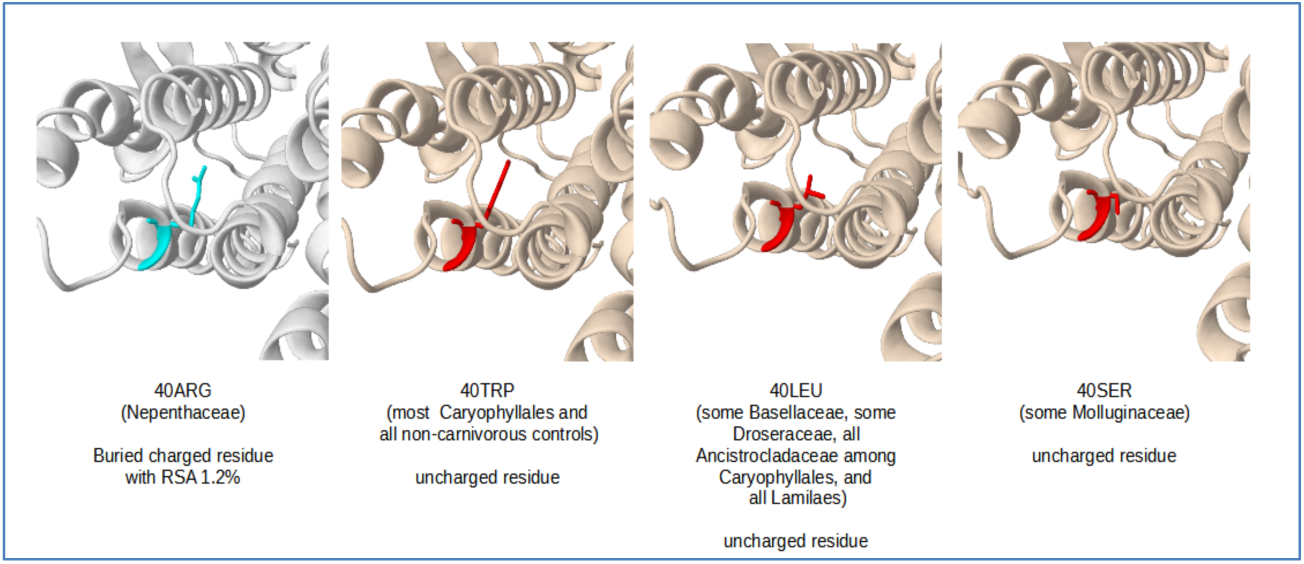
Characterization of structural impact of *Nepenthaceae*-specific amino acid changes in ndhD.

### Nepenthaceae-specific changes in psaA and ndhD across other carnivorous plants

The specificity of observed structure-impacting 147GLY and 40ARG changes in *psaA* and ndhD, respectively to *Nepenthaceae* group only were confirmed after extending the comparison with four other orders within the carnivorous plants and 11 non-carnivorous plant species, for which *psaA* and *ndhD* protein sequences were present in the NCBI database. These included 40 *psaA* and 16 *ndhD* sequences (**Appendix S3**). In *psaA*, residue 147 is an ALA in all the 40 additional sequences, thus confirming the 147GLY change unique to *Nepenthaceae*. In *ndhD*, residue 40 is a LEU in the Lamiales and TRP in the other non-carnivorous control sequences. This confirms the 40ARG change to be unique to *Nepenthaceae*.

### Impact of individual structure-impacting mutations on the interaction between psaA and ndhD

A frequency polygon comparison of the estimated distance between the structure-impacting residues of *psaA* and *ndhD* for 300 models of *Nepenthes* and other groups suggested a 2.67% greater frequency of non-*Nepenthes* models in the 43.4—51.3 range and a 5.33% greater frequency of Nepenthes models in 67.4—83.3 distance range. This suggested a marginal but specific change in the *psaA*—*ndhD* inter-residue distance distribution for the *Nepenthes* structures compared to the other groups. There are 29 *Nepenthes* models in the 43.4—51.3 distance range but 37 non-*Nepenthes* models in the same range, while there are only 79 non-*Nepenthes* models in the 67.4—83.3 distance range but 95 *Nepenthes* models in the same range (**Figure 3A; Table 2**). We observed that the minimum distance between the concerned residues of the interacting proteins, *psaA* and *ndhD*, was >6Å greater in *Nepenthes* than in the non-*Nepenthes* group: 25.9Å in *Nepenthes* and 19.4Å in the non-*Nepenthes*. Further, the overall conformations of interacting structures with least distances respectively in the *Nepenthes* and non-*Nepenthes* groups (25.9 Å and 19.4 Å) were vastly different from each other (**Figure 3B**). In both models, it can be seen that the structural conformation for *psaA* shows no or marginal difference but that of *ndhD* interacting with *psaA* is significantly changed -*ndhD* on the right side of *psaA* for *Nepenthes* and on top of *psaA* for non-*Nepenthes*.

**Figure 3.**
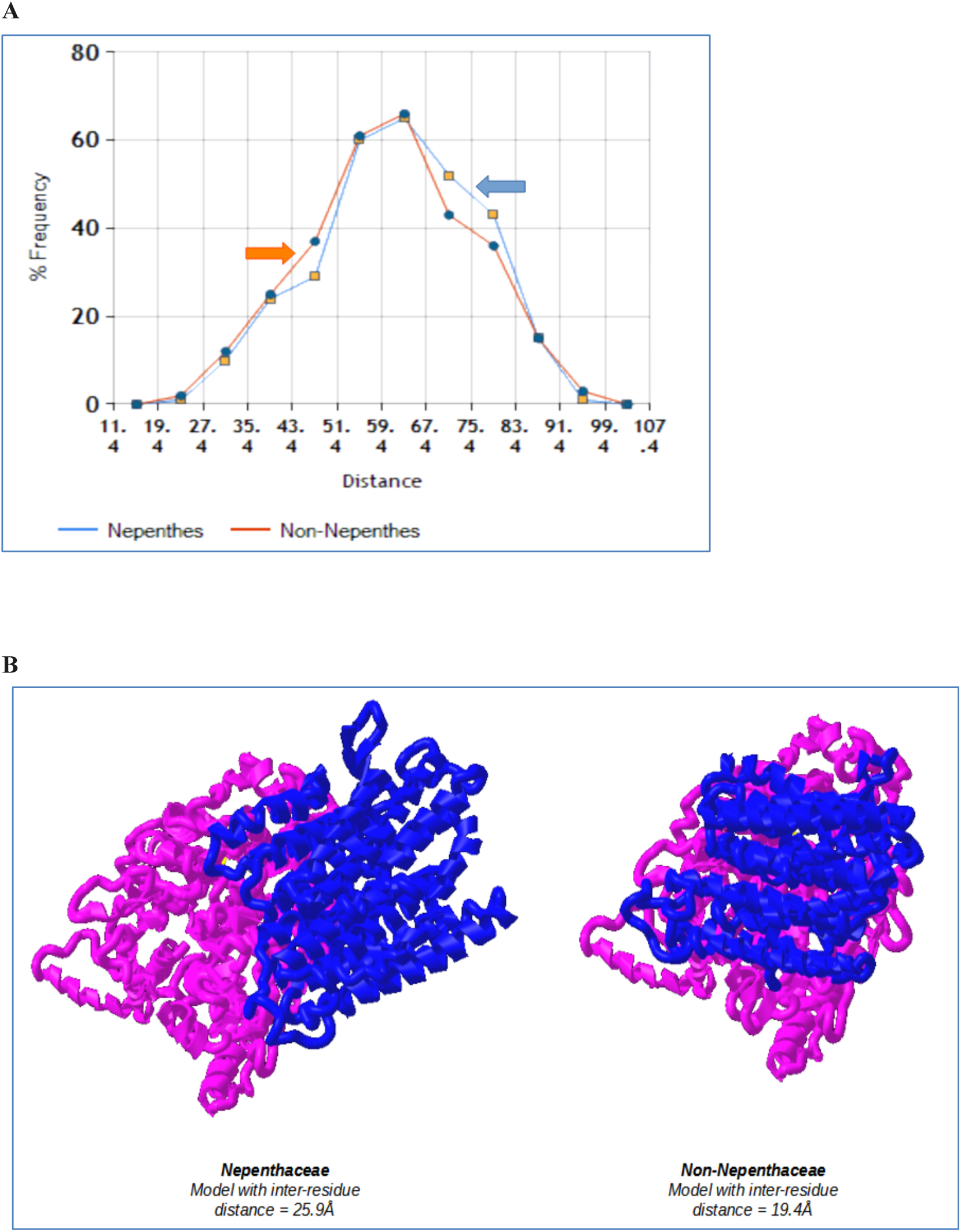
Frequency polygon of distance (Angstroms) between structure impacting residues 147 in psaA and 40 in ndhD, in *Nepenthes* and other groups in 300 models of interactions between the proteins. Classes with a distance range of 8 were formed with the lowest class value being 11.4 and highest-class value being 107.3 (A), 3D structural visualization of protein-protein interactions for the conformation with the least distance between the structure impacting residues 147 in psaA and 40 in ndhD (B).

**Table 2.** Frequency Table (A) and Polygon statistics (B) for analyses of the 300 predicted psaA-ndhD interaction models, each for *Nepenthes* and for the others.

| A |  |  | B |  |  |
| --- | --- | --- | --- | --- | --- |
| Frequency Table | Nepenthes | Others | Polygon Statistics - Group specific | Nepenthes | Others |
| Class | Count | Count | Lowest Distance | 25.9 | 19.4 |
| 11.4-19.3 | 0 | 0 | Highest Distance | 97.5 | 93.9 |
| 19.4-27.3 | 1 | 2 | Polygon Statistics - Combined | Nepenthes & Others |  |
| 27.4-35.3 | 10 | 12 | Total Number of Values | 300 |  |
| 35.4-43.3 | 24 | 25 | Number of Distinct Values | 231 |  |
| 43.4-51.3 | 29 | 37 | Lowest Class Value | 11.4 |  |
| 51.4-59.3 | 60 | 61 | Highest Class Value | 107.3 |  |
| 59.4-67.3 | 65 | 66 | Number of Classes | 12 |  |
| 67.4-75.3 | 52 | 43 | Class Range | 8 |  |
| 75.4-83.3 | 43 | 36 |  |  |  |
| 83.4-91.3 | 15 | 15 |  |  |  |
| 91.4-99.3 | 1 | 3 |  |  |  |
| 99.4-107.3 | 0 | 0 |  |  |  |

## DISCUSSION

### Complex adaptations in plant carnivory

Plant carnivory is a classic example of adaptation for plants to survive in dense nutrient-poor habitats, with wet or sunny and occasionally dry soil extremes (Adamec, 2002, Paniw et al., 2017). Studying these adaptations at a molecular level has been made possible by the advances in high throughput DNA sequencing (Aranguen-Diaz et al., 2018; Murphy et al 2020, Butts et al., 2016, Jensen et al., 2015, He et al., 2015). These adaptive molecular changes generate a novel function or modify an existing function in carnivorous plants, primarily via gene duplication and divergence. The adaptive functions are specific to metabolism and nutrient absorption, digestion of animal prey, the evolution of biomechanical factors such as formation, texture, regulation, maintenance, and movements of specific traps (Lan et al., 2017, Silva et al., 2019, Whitney and Federle, 2013), and energy optimization for trap activation (Laakkonen et al., 2006, Jobson et al., 2004). Metabolomics methods have been proven to facilitate carnivory (Hatcher et al., 2020, Hatcher et al., 2021). It has been suggested that there could be taxon-specific functional convergence of molecular changes behind these complex adaptations (Wheeler and Carstens, 2018).

### Nepenthes growth environment and trade-off between photosynthesis and carnivory

*Nepenthes* species are unique among the carnivorous plants in producing a pitcher from the tip of an elongated leaf. These plants grow in abundant light or water conditions but nutrient-poor soil. Their prey ranges from Formicidae, Hymenoptera, Coleoptera to Diptera (Chin et al., 2014), from which they absorb macronutrients such as nitrogen (Ellison, 2006, Moran et al., 2001, Schulze et al., 1997), phosphorus, potassium and magnesium (Adamec, 2002, Pavlovic et al., 2014). As per Givnish’s cost and benefit model (Givnish et al., 1984), due to the growing conditions, the benefits of capturing and feeding on prey exceed the costs of carnivory, such as the development of non-photosynthetic structures like specialized traps, digestive glands and enzymes. However, Givnish’s cost and benefit model is a conjecture at best because it fails to consider these plants’ compromised root absorption capacity as a benefit (Ellison, 2006, Wakefield et al., 2005). There is also the cost of energy needed for the respiration of these carnivorous organs. Feeding on the prey is known to increase the efficiency of photosynthesis in carnivorous plants (Capo-Bauca et al., 2020, Pavlovic et al., 2014), although conflicting observations have been made (Pavlovic et al., 2010, Wakefield et al., 2005). Recent studies also provide some evidence of spatio-temporal changes in photosynthesis and respiration at the time of prey capture (Pavlovic, 2010).

### What does cyclic electron transport do and what could this adaptation mean?

Evidence suggests that during prey retention, there might be a temporary slowing of photosynthesis to stimulate the carnivorous organs’ respiration (Pavlovic et al., 2010), and a transient switch mediates this regulation from the linear to cyclic electron transport process (Maurer et al., 2020) for achieving ATP homeostasis. In cyclic electron transport, the photosystem I (PSI) subcomplex plays a key role (Yamori et al., 2016) by forming a super-complex with the NDH sub-complex (Shen et al., 2021) along with other helper genes. The PSI-NDH super complex stabilizes ndh, especially in high-stress conditions (Shikanai, 2016). Although discovered based on their homology with other genes in the respiratory complex, the chloroplast-encoded *ndh* genes play a vital role as part of the photosynthetic machinery by regulating the ATP:NADPH ratio to offer photoprotection in strong light conditions (Shikanai, 2016).

### *Nepenthaceae-specific amino acid changes in* psaA *and* ndhD

Our study found amino acid changes, one in *psaA* and one in *ndhD*, specific to *Nepenthaceae* that impacted their respective three-dimensional structures. Given that photosystem I and ndh subunits interact to form a super-complex, we tested the effect of these structure-impacting amino acid changes in *psaA* and *ndhD* on the structural interaction. We found that the interaction was also impacted (**Figures 1-3**). These observations suggest *Nepenthaceae*-specific changes in the psI-ndh super complex may aid the unique nature of the heterotrophic carnivory observed in these species. This is only a conjecture at present and can be confirmed by sampling across more *Nepenthes* species and sequencing their chloroplast genomes, and by modeling a complete super complex including the helper genes.

## CONCLUSION

We observed *Nepenthes*-specific amino acid changes in the protein psaA and ndhD that impact their three-dimensional structures and the interactions between them. This may be responsible for *Nepenthaceae*-specific adaptations in the photosynthetic machinery involving cyclic electron transport process.

